# Single Cell Analysis Reveals Partial Reactivation of X-chromosome Instead of Chromosome-wide Dampening in Naïve Human Pluripotent Stem Cells

**DOI:** 10.1101/697565

**Authors:** S Mandal, D Chandel, H Kaur, S Majumdar, M Arava, S Gayen

## Abstract

Recently, a unique form of X-chromosome dosage compensation has been demonstrated in human preimplantation embryos, which happens through the dampening of X-linked gene expression from both X-chromosomes. Subsequently, X-chromosome dampening has also been demonstrated in female human pluripotent stem cells (hPSCs) during the transition from primed to naïve state. However, the existence of dampened X-chromosomes remains controversial in both embryos and hPSCs. Specifically, in preimplantation embryos it has been shown that there is inactivation of X-chromosome instead of dampening. Here, we have performed allelic analysis of X-linked genes at the single cell level in hPSCs and found that there is partial reactivation of the inactive X-chromosome instead of chromosome-wide dampening upon conversion from primed to naïve state. In addition, our analysis suggests that the reduced X-linked gene expression in naïve hPSCs might be the consequence of erasure of active X-chromosome upregulation.

## Introduction

In therian mammals, to balance the X-chromosome dosage between males and females, one X-chromosome becomes inactivated in female cells (Lyon, 1961). The dosage imbalance between a single active X-chromosome and two copies of autosomes (AA) is compensated through upregulation of the active-X chromosome in both males and females (Deng et al., 2011; Larsson et al., 2019; Ohno S, 1967). Recently, another form of X-chromosome dosage compensation has been demonstrated in human preimplantation embryos, termed as X-chromosome dampening. Based on single cell transcriptome analysis of human preimplantation embryos, Petropoulos *et al.* (2016) found that X-linked gene expression gradually decreased from morula to blastocyst stage, while both X-chromosomes were maintaining active state (Petropoulos et al., 2016). Based on these, they proposed that dampening of X-linked gene expression from both X-chromosome as a likely dosage compensation mechanism during human pre-implantation development. However, dampening phenomenon in human embryos remains controversial (De Mello et al., 2017; Saiba et al., 2018). De Mello *et al.* (2017) found evidence of inactivation of the X-chromosome instead of dampening in preimplantation embryos when they reanalyzed the same transcriptome dataset of Petropoulos *et al.* (2016) with more stringency. In addition, Sahakyan *et al.* (2017) showed that naïve human pluripotent stem cells (hPSCs) also exhibit the X-chromosome dampening found in embryos (Sahakyan et al., 2017a). However, similar to the human embryos, X-chromosome states in hPSCs remains unclear (Kaur et al., 2019; De Mello et al., 2017). Conventional hPSCs derived from blastocysts represent a primed state instead of naïve state and are therefore unable to recapitulate the preimplantation X-chromosome states (Nichols and Smith, 2009; Sahakyan et al., 2017b). To model the preimplantation X-chromosome state, Sahakyan *et al.* (2017) converted primed hPSCs to the naïve state using 5iLAF culture condition (Fig. 1A) (Theunissen et al., 2014). The primed cell line used for their study, UCLA1, harbored one active-X chromosome and one inactive-X chromosome. The transition of primed to naïve state happened through an intermediate early naïve state (Fig. 1A). Primarily based on RNA-sequencing analysis using bulk cell population, they suggested that the inactive X-chromosome was reactivated upon transition from primed to the early naïve state and this was followed by X-chromosome dampening in late naïve cells (Fig. 1A). However, considering the heterogeneity of cell states during the conversion process, in this study, we have analyzed available single cell RNA-Seq (scRNA-Seq) dataset of early and late naïve cells from Sahakyan *et al.* (Sahakyan et al., 2017a), to gain better insight into X-chromosomal states (Fig. 1A).

**Figure 1:**
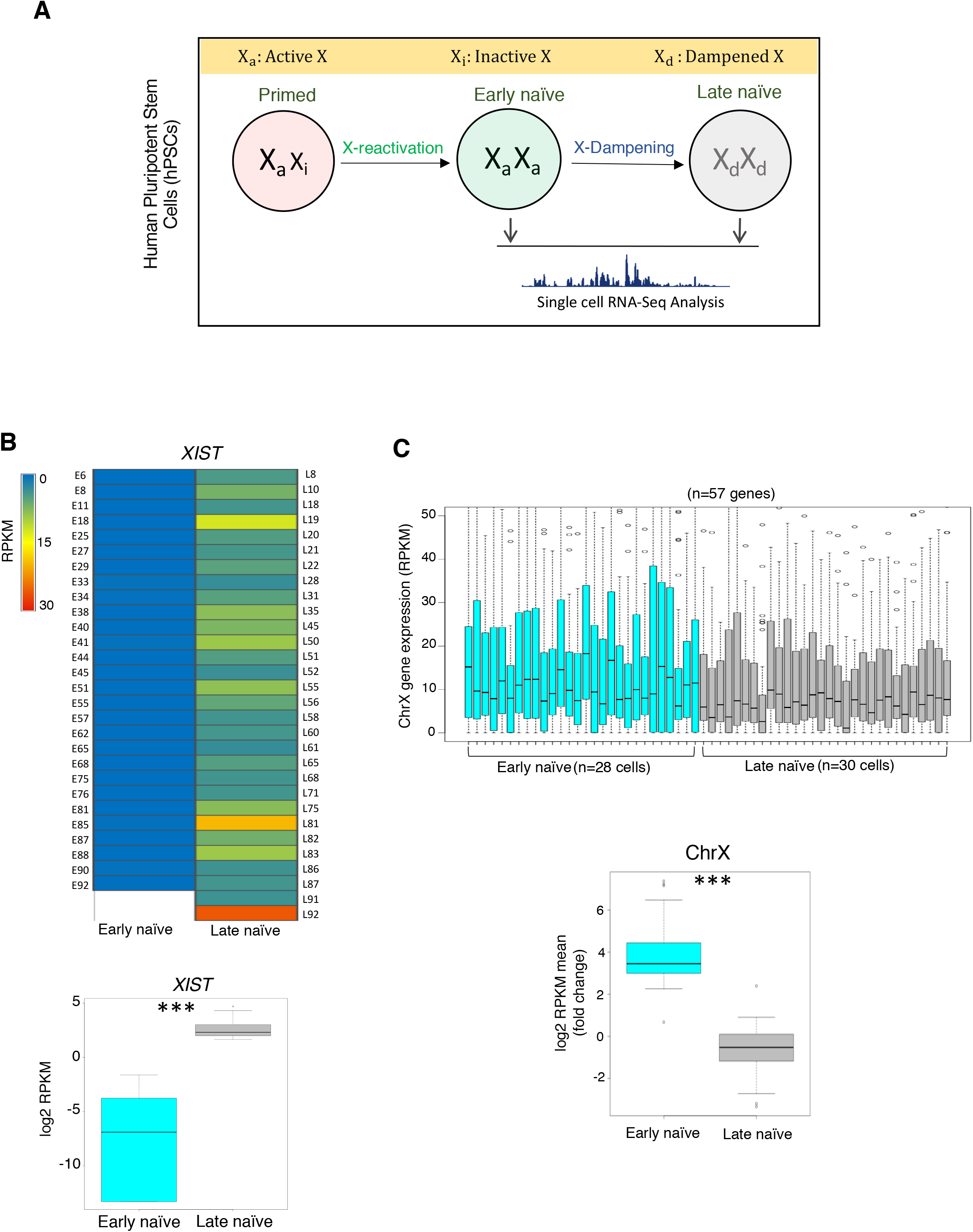
Transition of early to late naïve state is associated with increased *XIST* expression and reduction in X-linked gene expression. (A) Schematic representation of different stages and corresponding X-chromosome states of conversion of primed hPSCs to naïve state as described in Sahakyan *et al.* (2017). In this study, we performed analysis of scRNA-Seq dataset acquired from the early and late naïve state. (B) Comparison of *XIST* expression (RPKM) between early naïve (n=28 cells) and late naïve cells (n=30 cells). *p*<0.00001 (Mann-Whitney U-test) (C) Comparison of X-linked gene expression (n=57 genes) between early and late naïve cells. *p*< 0.00001 (Mann-Whitney U-test).

## Results

### Increased *XIST* expression and reduction in X-linked gene expression upon conversion of early to late naïve state

First, in early and late naïve cells, we quantified the expression of *XIST,* a master regulator of X-inactivation. We found that a majority of the early naïve cells had very low level of *XIST* expression, whereas late naïve cells mostly showed higher level of *XIST* expression (Fig. S1A). Overall our analysis revealed that transition of early to late naïve state was associated with significant increase of *XIST* expression (Fig. S1B). Next, based on quality (RPKM sum and mean) and *XIST* expression level, we selected the top 28 early cells having low level of *XIST* and the top 30 late cells having higher level of *XIST* expression for further analysis (Fig. 1B). Again, comparison of *XIST* expression in these cells (28 early vs 30 late) showed a significant increase in late naïve cells compared to early naïve cells (Fig. 1B). In addition, we found that there was a significant reduction in X-linked gene expression in late naïve cells compared to early naïve cells (Fig. 1C; Supplementary file1). Altogether, our analysis of scRNA-Seq data showed increased *XIST* expression and reduction in X-linked gene expression upon transition from the early to late naïve state.

### Reduction in X-linked gene expression in late naïve cells is independent of *XIST*

It has been shown that X-chromosome dampening in preimplantation embryos is associated with the expression of *XIST* from both X-chromosomes (Petropoulos et al., 2016). In fact, X-dampening initiates concomitantly with the initiation of *XIST* expression and therefore it is thought that *XIST* might have an important role in the dampening process. To test this, we examined what fraction of cells show *XIST* expression from both X-chromosomes in late naïve cells and if the reduction in X-linked gene expression is restricted to the biallelically *XIST* expressed cells. We found about 26*%* cells (9 of 35) expressed *XIST* from both X-chromosomes (Fig. 2A; Supplementary file 2). Interestingly, comparison of the global X-linked gene expression level of *XIST* biallelic vs monoallelic cells did not show any significant difference (Fig. 2B; Supplementary file 2). Moreover, we observed that there was no significant difference in X-linked gene expression when we compared against the *XIST* negative cells (Fig. 2B; Supplementary file 2). Based on these data, we concluded that reduction in X-linked gene expression in late naïve cells was independent of *XIST*.

**Figure 2:**
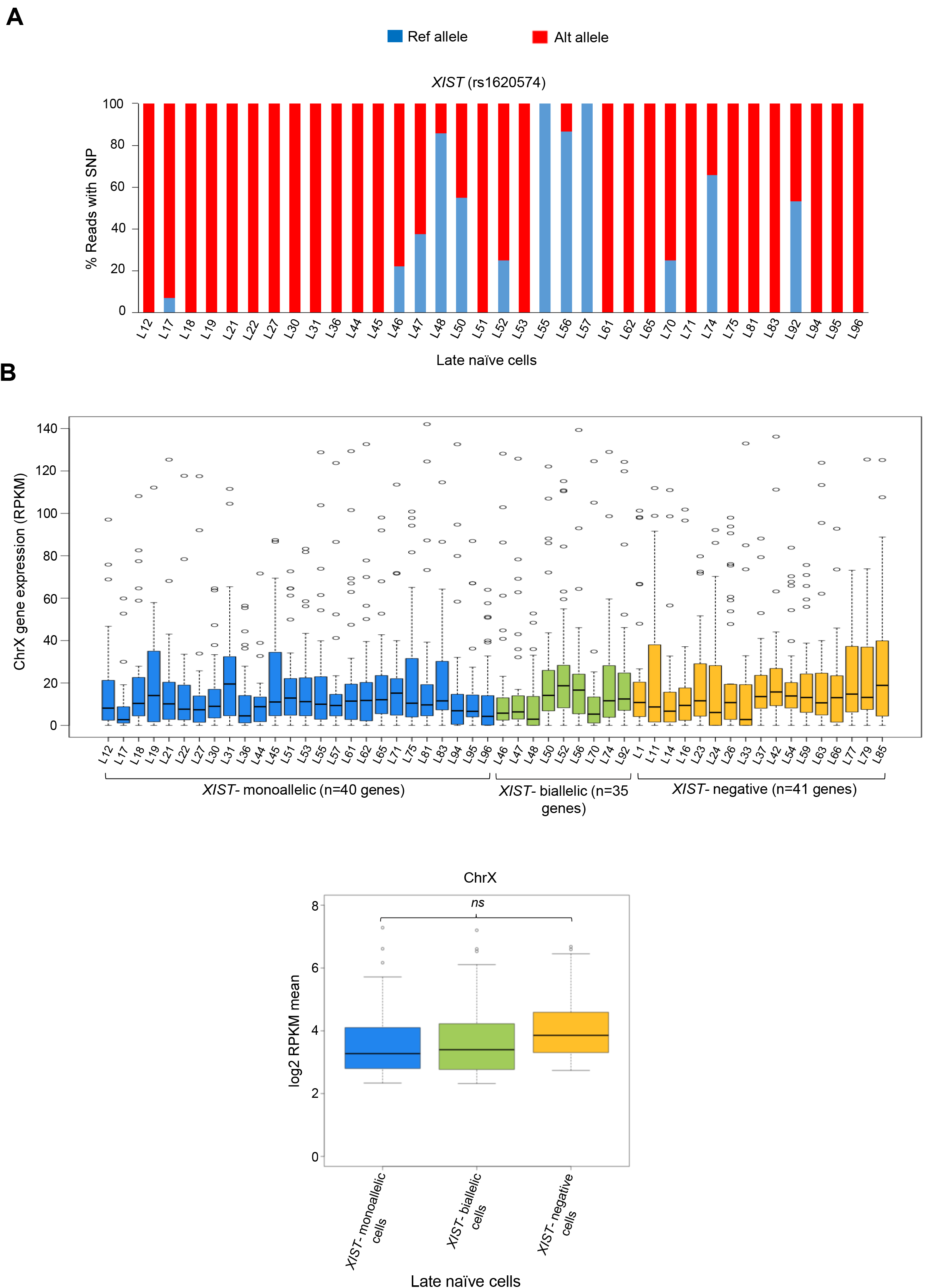
Reduction in X-linked gene expression in late naïve cells is independent of *XIST*. (A) Allelic expression of *XIST* in late naïve cells (n=35 cells). (B) Comparison of X-linked gene expression among *XIST* -monoallelic, *-*biallelic and -negative cells. Non-significant at *p*<0.05 (Mann Whitney U-test).

### No evidence of X-chromosome inactivation or dampening

We then looked into possible mechanisms behind the reduction in X-linked gene expression in late naïve hPSCs. First, we postulated that it could be due to the either X-chromosome inactivation or dampening. To test for X-inactivation, we profiled allelic expression of multiple X-linked genes distributed across the X-chromosome based on single nucleotide polymorphisms (SNPs) in early and late cells (Fig. 3; Supplementary file 3). We found that the majority of early cells showed monoallelic expression of many genes along with biallelically expressed X-linked genes, which indicated incomplete reactivation of X-linked genes (Fig.3A & 3B). Moreover, we found significant variation in the proportion of biallelic vs monoallelic genes between these cells (Fig. 3B). Interestingly, late naïve cells showed a similar pattern of allelic expression as early cells. However, there was a significant increase in the fraction of SNPs showing biallelic expression in late naïve cells compared to early naïve cells. (Fig. 3B). Altogether, these data indicated that late naïve cells do not harbor inactive X-chromosome but rather they have partially reactivated X-chromosome. If the late naïve cells underwent X-inactivation, then monoallelic expression of most of the X-linked genes would be expected. In contrast, we observed increased biallelic expression upon transition from early to late stage. Next, we examined whether X-dampening was causing reduction of X-linked gene expression in late naïve cells. From the allelic analysis of X-linked gene expression it was clear that late naïve cells harbor partially reactivated X-chromosome. If these cells harbored dampened X-chromosomes, we would have expected biallelic expression of most of the genes chromosome-wide, which was not observed. In addition, we compared median expression of biallelically expressed genes of early cells to that of the late naïve cells. If X-dampening was occurring then we would have expected a significant decrease in median expression of biallelically expressed genes in late cells. However, significant differences were not observed (Fig. 3C). Taken together, we concluded that there was neither X-inactivation nor dampening upon conversion of early to late naïve cells. Second, to determine whether loss of an X-chromosome in late cells is causing the reduction in X-linked gene expression in late naïve cells, we explored X-chromosome ploidy of these cells. It was clear that cells harbored two X-chromosomes as evident by the biallelic expression of some X-linked genes (Fig. 3A). However, the possibility existed that these cells may lose part (s) of the X-chromosome. To test for this, the gene expression ratio of X-linked genes across the X-chromosome was analyzed, but significant differences between early vs late naïve cells were not identified (Fig. 3D; Supplementary file 4). Therefore, we confirmed that loss of a portion of the X-chromosome is not the cause of reduction in X-linked gene expression in late naïve cells.

**Figure 3:**
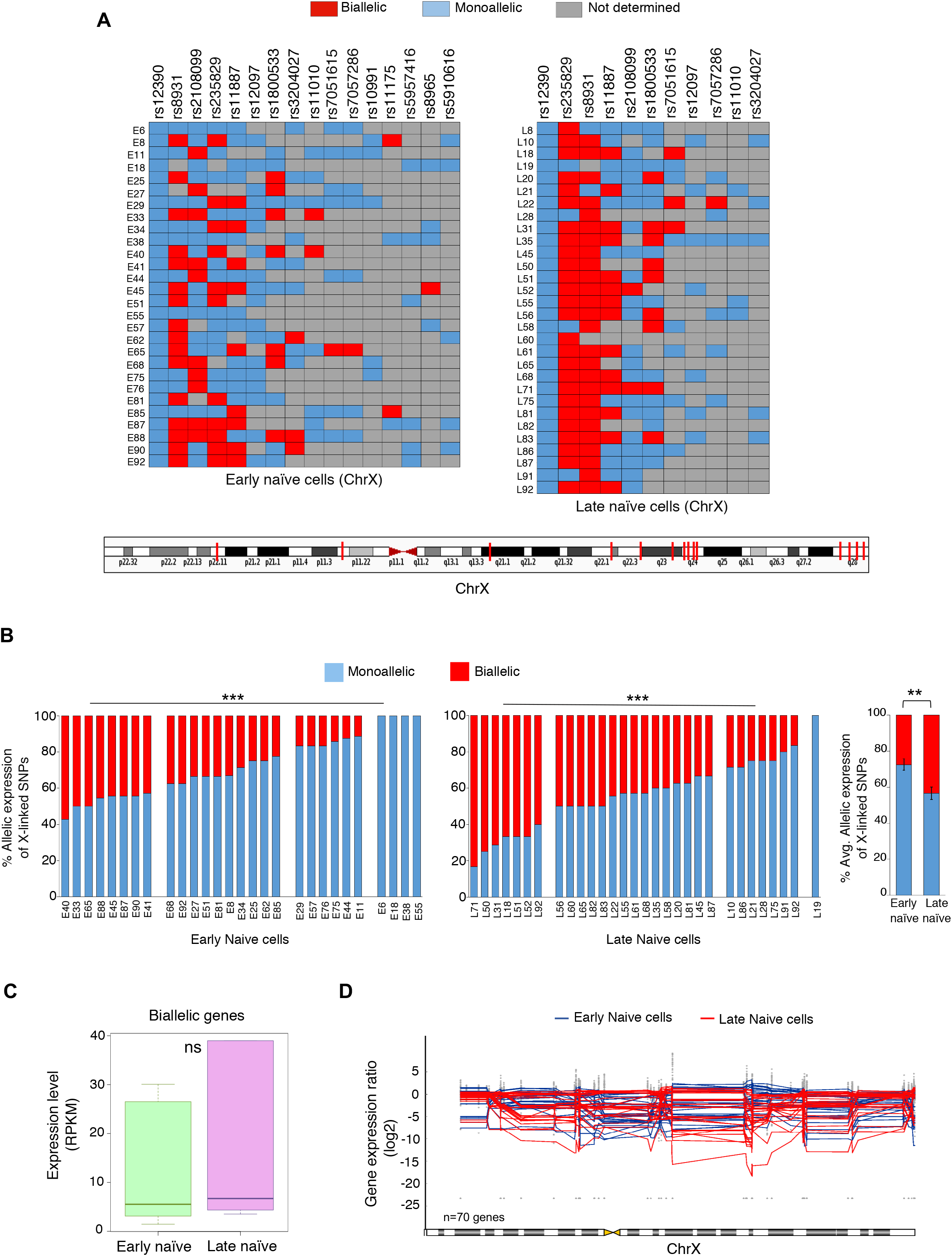
Partial reactivation of inactive-X chromosome upon conversion of primed to naïve state. (A) Allelic expression analysis of X-linked genes in early and late naïve cells at single cell level. Bottom, genomic position on the X-chromosome of the X-linked genes analyzed (B) Histogram showing the percent of SNPs showing monoallelic and biallelic in each cell of early and late naïve state. *p*<0.00001, *p*<0.01(Avg. plot) (Student’s t-test) (C) Comparison of median of expression level of biallelically expressed genes between early and late naïve cells. Non-significant at *p*<0.05 (Mann Whitney U-test) (D) Ploidy analysis of the cells of early and late naïve state through the analysis of gene expression ratio across the X-chromosome.

### Erasure of active-X upregulation might be the cause of reduced X-linked gene expression in late naïve cells

Next, we investigated if the erasure of active X-chromosome upregulation might be causing the reduction in X-linked gene expression in late naïve cells as proposed by De Mello *et al. (De* Mello et al., 2017). In recent years, the existence of upregulated active X-chromosome has been extensively demonstrated in mammals as hypothesized by Ohno (Deng et al., 2011, 2013; Li et al., 2017; Lin et al., 2011; De Mello et al., 2017; Sangrithi et al., 2017). Although some studies have found lack of active-X upregulation (Chen and Zhang, 2016; Xiong et al., 2010). To probe this further, we analyzed X to autosomal (X:A) gene expression ratio of 7 different male primed hPSC lines. If a diploid male cell has upregulated active-X and then the X:A ratio should be more than 0.5 and closer to 1. Indeed, we found the X:A ratio of all male primed cells was greater than 1, indicating that primed hPSCs harbor an upregulated active X-chromosome (Fig. 4A; Supplementary file 5). We then asked whether the active-X upregulation becomes erased in naïve hPSCs. To test this, we compared the X:A ratio of male primed cells against different male naïve cell lines. Interestingly, a significant reduction of X:A ratio in naïve cells compared to the primed cells was observed, suggesting erasure of active X-chromosome upregulation in naïve cells (Fig. 4A). We made sure that the X:A ratio was not impacted by the difference between X-linked and autosomal gene expression distribution for each dataset (Fig. 4B). We focused on male cells for analysis of active-X upregulation as in female cells X-linked gene expression is often confounded with X-chromosome inactivation / reactivation / erosion. However, we profiled the X:A ratio in 3 different primed female cells (including UCLA1), which are known to harbor one inactive-X chromosome (Fig. 4C). We found the X:A ratio of female primed cells was above 1, which indicated that these cells harbor an upregulated active-X chromosome (Fig. 4C; Fig. S2B). Next, we examined the X:A ratio dynamics during the primed to naïve conversion of UCLA1 female cells. An significant increase in the X:A ratio upon transition from primed to early naïve state was observed (Fig. 4D; Fig. S2B). Considering the partial X-reactivation upon transition of primed to early naïve cells, it is obvious that the X:A ratio should increase provided that the active-X upregulation is not completely erased. If there was complete erasure of active-X upregulation, the X:A ratio should not have an observable increase. Therefore, this data indicated that erasure of active-X upregulation was incomplete in the early naïve state. Conversely, a decrease in the X:A ratio was observed in the late naïve state compared to the early naïve cells, which suggested that erasure of active-X upregulation was occurring (Fig. 4D; Fig. S2B). In this scenario, it is possible that the erasure of active-X upregulation led to the overall reduction in X-linked gene expression in late naïve cells. However, the decrease in X:A ratio from early to late naïve state was not significant as expected. We think that a significant increase in biallelic X-linked gene expression from early to late naïve state was masking this to some extent. In summary, our data suggests that X-dampening is not the determining factor for the reduction in X-linked gene expression in late naïve cells, and erasure of active-X upregulation might be leading to the decrease of X-linked gene expression (Fig. 4E).

**Figure 4:**
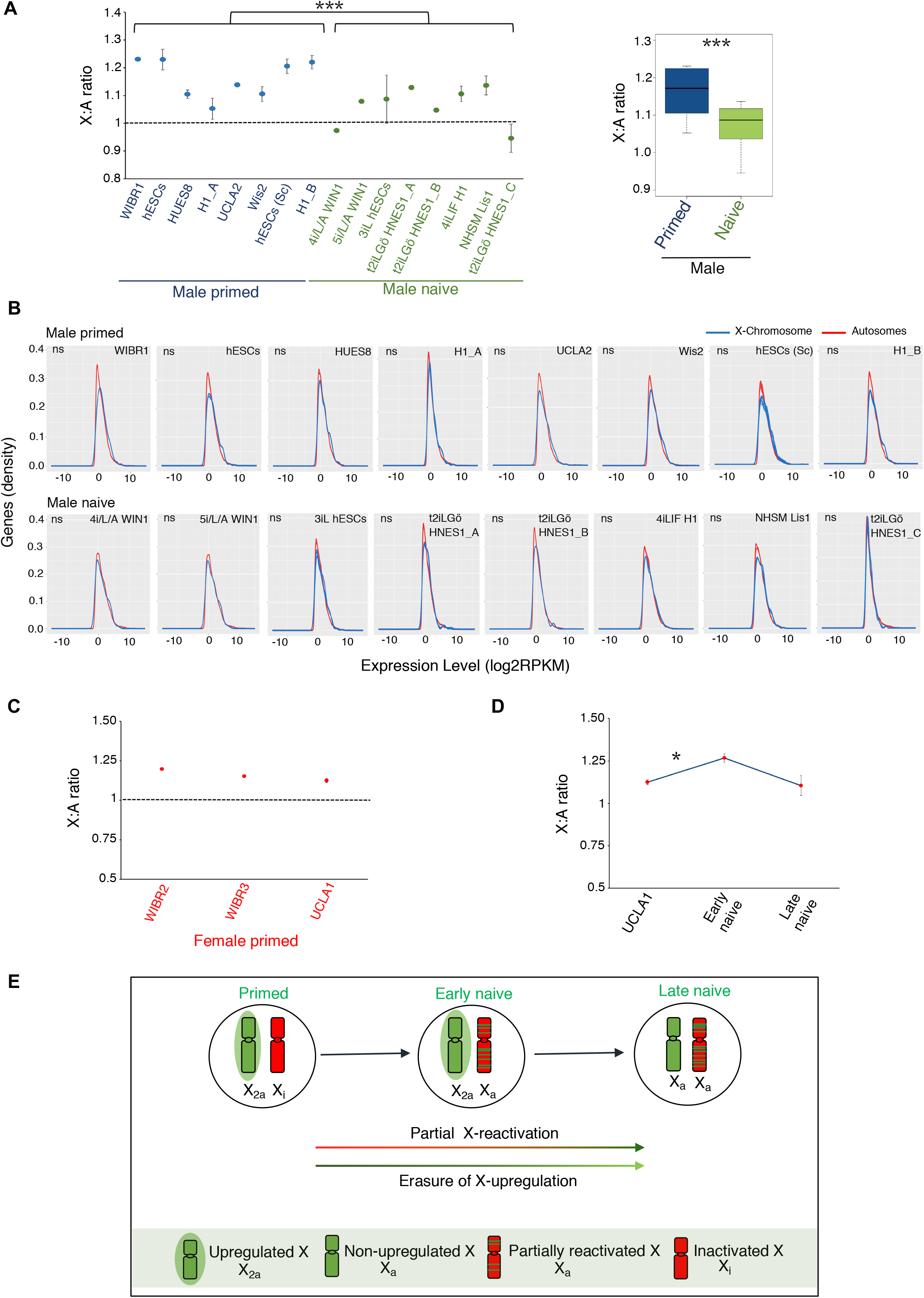
Reduction in X-linked gene expression in naïve hPSCs might be due to the erasure of active-X upregulation. (A) Comparison of X:A ratio between male primed and naïve hPSCs. *p*<0.001(Student’s t-test). (B) Histograms representing the distribution of X-linked and autosomal gene expression for male primed and naïve hPSCs with their different replicates (p > 0.05, by Kolmogorov-Smirnov test). (C) Analysis of X:A ratio in different primed female hPSCs. (D) Comparison of X:A ratio in UCLA1 primed, early naïve (cl4) and late naïve cells (cl9 & cl12). *p*<0.05 (E) Proposed model representing the X-chromosome states during the conversion of primed hPSCs to the naïve state.

## Discussion

X-chromosome states in female human preimplantation embryos remains elusive till date. While it has been reported that preimplantation embryos carry dampened X-chromosomes, other studies have provided evidence of inactivation of one of the X-chromosomes. One of the major challenges to resolve this issue is the lack of availability of surplus number of human embryos for experimentation. Therefore, hPSCs derived from human embryos serve as an alternative system. However, conventional hPSCs represent the primed state instead of the naïve state of preimplantation embryos (Davidson et al., 2015; Sahakyan et al., 2017b). Recently, Sahakyan *et al.* (2017) converted primed hPSCs to naïve state to model the X-chromosome states of preimplantation embryos and suggested that naïve hPSCs also carry dampened X-chromosomes. However, our analysis indicates that the conversion of primed hPSCs to naïve state is associated with the partial reactivation of the inactive-X chromosome instead of the chromosome-wide dampening (Fig. 4E). The main reason behind the dissimilar outcomes between our study and Sahakyan *et al.* (2017) is likely that their conclusion is primarily based on analysis of bulk RNA-sequencing of cell population, whereas our conclusion is based on analysis of single cell RNA-Seq dataset. Since, single cell RNA-Seq provides better clarity on cellular state and gene expression to distinguish the heterogeneity among cells in a population, we believe that our analysis provides better insight into X-chromosome states of hPSCs. In addition, our study suggests that erasure of active-X upregulation might be causing the reduction in X-linked gene expression in late naïve cells (Fig. 4E).

Although two studies led to the dissimilar outcomes, several observations in our analyses were consistent with findings by Sahakyan *et al.* (2017). For example, we also found that early naïve cells were mostly *XIST* negative and transitioned to *XIST* positive in the late naïve state which was accompanied by a reduction in X-linked gene expression (Fig. 1). Moreover, about 26% of cells in late naïve state showed *XIST* expression from both X-chromosomes, which was similar to what was reported by Sahakyan *et al.* (2017). We should point out that in preimplantation embryos majority of the cells (~85%) express *XIST* from both X-chromosomes (Okamoto et al., 2011) and it is believed that *XIST* might have an important role in X-dampening. However, we found that reduction in X-linked gene expression was independent of *XIST* as *XIST-*biallelic, -monoallelic and -negative cells showed almost similar level of gene expression in late naïve cells (Fig. 2B).

On the other hand, allelic analysis of X-linked gene expression at the single cell level revealed striking differences between the two studies. We found that early and late naïve cells harbored partially reactivated X-chromosome as indicated by monoallelic expression of many genes in each cell of both cell states (Fig. 3). This was contrary to Sahakyan *et al.* (2017) where they found most of the genes had biallelic expression. Importantly, we found significant variations in allelic patterns of gene expression in different cells within a population, such as same genes, which were monoallelic in some cells, showed biallelic expression in other cells. In this scenario, bulk cell population analysis will always show biallelic gene expression, which was might be the case for Sahakyan’s observation. Interestingly, while Sahakyan *et al.* (2017) interpreted the reduction in X-linked gene expression in late naïve cells as dampening, we found lack of dampening since there was no significant difference in the median expression of biallelically expressed genes between early and late naïve cells (Fig. 3C). In fact, Theunissen *et al.* (2016) found that female naive cells had significantly higher X-linked gene expression compared to male naive cells, which also suggested that naïve female cells harbored active X-chromosomes instead of dampened X-chromosomes (Theunissen et al., 2016). In addition, we observed that the proportion of biallelically expressed genes increased significantly in late naïve cells compared to early naïve cells, which indicated that the cells were still undergoing the reactivation process.

Some observations by Sahakyan *et al.* also indicated that there was an incomplete erasure of epigenetic memory on the inactive X in the naïve state. They found, upon differentiation, naïve hPSCs underwent non-random X-inactivation dissimilar to the normal development, and the same X-chromosome that was inactive in the primed hPSCs was again inactivated. Even, naïve cells showed an accumulation of H3K27me3 repressive marks on one of the X-chromosomes, that had not been observed in preimplantation blastocysts (Okamoto et al., 2011). We think the presence of H3k27me3 is the result of incomplete erasure of inactive-X marks, which is consistent with our observation of a partially reactivated X-chromosome. Taken together, these findings suggest that naïve cells were still in the process of removing inactive-X epigenetic marks.

Our study suggests that erasure of active-X upregulation might be causing the reduction in X-linked gene expression upon transition from early to late naïve state. Many recent studies have demonstrated the existence of upregulated active-X chromosomes in mammals (Deng et al., 2013; Larsson et al., 2019). We also found evidence for upregulated active-X in 7 different male primed hPSC lines (Fig. 4A). Importantly, our analysis of male primed and naïve cells strongly indicates that there is erasure of active-X upregulation upon conversion of primed to naïve state (Fig. 4A). Erasure of upregulation has been demonstrated previously in spermatids, during oogenesis and germ cell reprogramming of both sexes (Di and Disteche, 2006; De Mello et al., 2017; Sangrithi et al., 2017). Moreover, our observation is consistent with some other studies in mouse, which showed that naïve ESC has lower X:A ratio compared to the differentiated cells (Lin et al., 2007; Marks et al., 2015). Specially, during female ESC differentiation, upregulation increases concomitantly with the initiation of X-inactivation (Larsson et al., 2019). In fact, Theunissen *et al.* (2016) found a reduction in X-linked gene expression in male naïve hPSCs compared to that in primed, which may be due to the erasure of active X-chromosome upregulation (Theunissen et al., 2016). Furthermore, it has been shown that male blastocyst had significantly lower X:A ratio compared to the primed hPSCs (De Mello et al., 2017). Taken together, we believe that primed to naïve conversion of hPSCs is accompanied by erasure of active-X upregulation. In addition, our data indicate that the erasure of upregulation for female UCLA1 is might still ongoing in the early naïve state and only reaches near the completion in the late naïve state, thereby leading to the reduction in X-linked gene expression upon transition from early to late naïve state (Fig. 4D). In fact, during germ cell reprogramming it has been shown that erasure of upregulation and reactivation of inactive-X does not occur simultaneously rather loss of upregulation occurs later than loss of X-inactivation (Sangrithi et al., 2017).

Collectively, our study indicates that the conversion of primed to naïve state is associated with the incomplete reactivation of X-chromosome rather than X-inactivation or X-dampening. Importantly, our data also indicates that erasure of active-X upregulation might be leading the reduction in X-linked gene expression in naïve hPSCs. Although our results argue against dampening and propose erasure of active-X upregulation is leading to the reduction in X-linked gene expression in late naïve cells, further work must be done with better scRNA-Seq data. Finally, better culture conditions are necessary to establish naïve hPSCs that recapitulate the X-chromosome states of preimplantation embryos.

## Experimental procedures

### Data acquisition

RNA-Seq datasets were acquired from Gene Expression Omnibus (GEO) under the accession number GSE87239 (Sahakyan et al., 2017a). For additional datasets see the supplementary experimental procedures.

### Variant calling

First, reads were mapped to the human genome (hg38) using STAR. To mark the duplicate reads from the aligned reads of single cells, we used Picard tools v2.18.11 (https://broadinstitute.github.io/picard/). Next, we retrieved the allelic read counts for SNPs by using GATK (v3.8) “HaplotypeCaller”. We considered those SNPs for our analysis, which were present in UCLA1 cell line database (GSM2420529). Further we annotated those SNPs using dbSNP Build 152 (GRCh38.p12).

### Allelic expression analyses

For allelic expression analysis, we considered the SNPs having ≥3 reads per SNP site in a cell. Further, we proceeded with those SNPs having informative reads in at least five different cells of each category; early and late naïve. The allelic expression was calculated by directly counting the allele-specific reads covering a SNP position mapped to the reference or the alternative allele and then dividing it by the total number of reads covering that position. A SNP was considered monoallelic if at least 90% of the allelic reads was coming from only one allele. We only considered allelic ratios of SNPs for those genes, which had RPKM ≥ 1. Finally, we considered only those SNPs for which the allelic data was available in at least four cells for each early and late naïve cells. We validated allele specific expression pipeline through analysis of genes of an autosome (Chr17), which showed mostly biallelic expression of SNPs (Fig. S2A). Moreover, SNPs belongs to the same gene showed almost similar allelic expression pattern in most of the cells, except for few cells.

## Author’s Contribution

SG, SM (Susmita Mandal), DC, and HK conceptualized the study. SG supervised the study. Bioinformatic analyses was done by SM (Susmita Mandal) and DC. SG, SM (Susmita Mandal), DC, MA, and SM wrote, edited and proofread the manuscript. Final manuscript was edited and approved by all the authors.

## Acknowledgments

We thank Pavithra RV, Karunyaa M and Amitesh Panda for their help in art work and discussion. Study is supported by DBT grant (BT/PR30399/BRB/10/1746/2018), DBT-Ramalingaswamy fellowship (BT/RLF/Re-entry/05/2016) and IISC-MHRD Start up grant awarded to SG. We also thank DST-FIST [SR/FST/LS11-036/2014(C)], UGC-SAP [F.4.13/2018/DRS-III (SAP-II)] and DBT-IISc Partnership Phase-II (BT/PR27952- INF/22/212/2018) for financial support.

