## Supplementary figure and text for "Single Cell Analysis Reveals Partial Reactivation of X-chromosome Instead of Chromosome-wide Dampening in Naïve Human Pluripotent Stem Cells"

Fig. S1

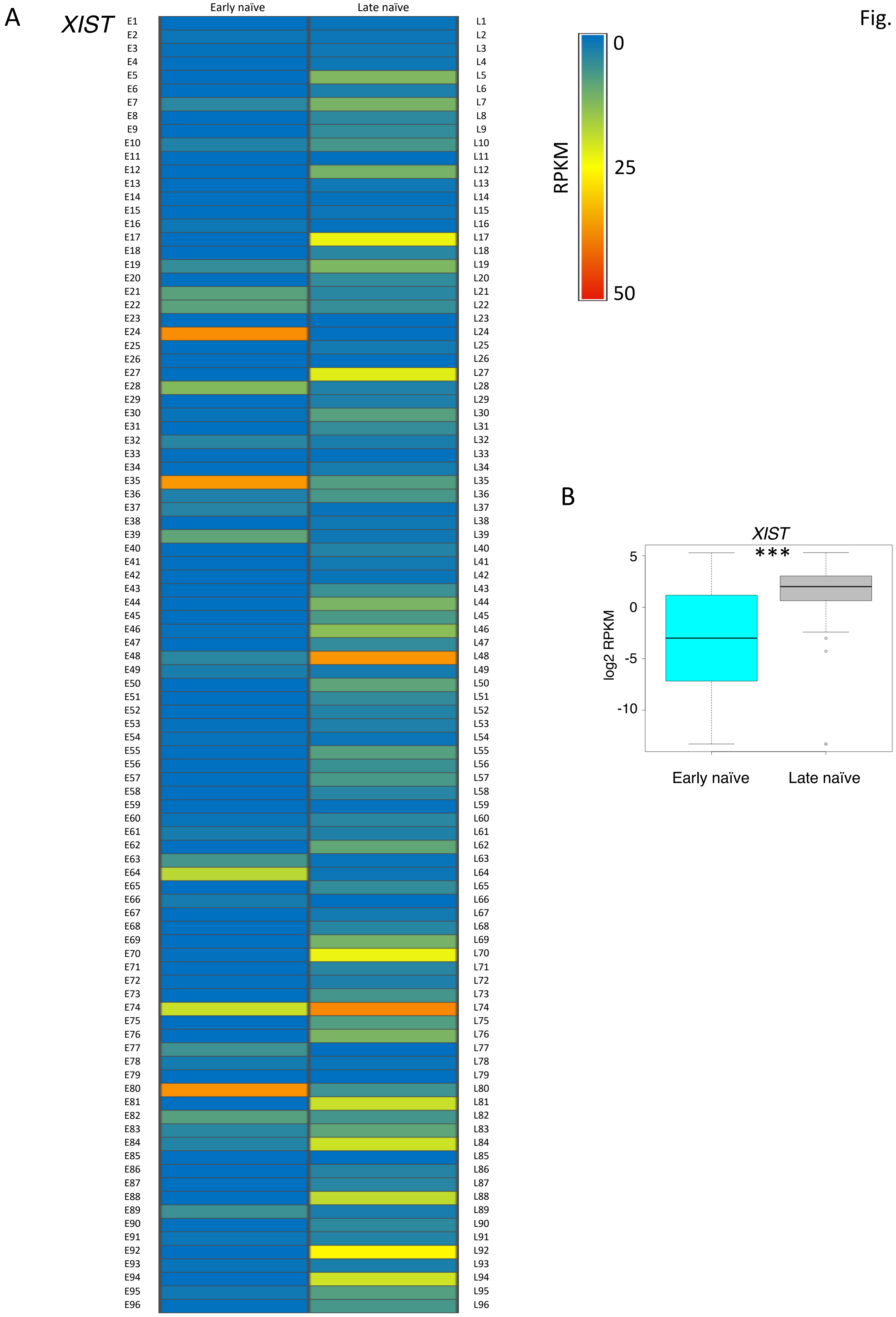
A
XIST
Early naïve
Late naïve
E1
E2
E3
E4
E5
E6
E7
E8
E9
E10
E11
E12
E13
E14
E15
E16
E17
E18
E19
E20
E21
E22
E23
E24
E25
E26
E27
E28
E29
E30
E31
E32
E33
E34
E35
E36
E37
E38
E39
E40
E41
E42
E43
E44
E45
E46
E47
E48
E49
E50
E51
E52
E53
E54
E55
E56
E57
E58
E59
E60
E61
E62
E63
E64
E65
E66
E67
E68
E69
E70
E71
E72
E73
E74
E75
E76
E77
E78
E79
E80
E81
E82
E83
E84
E85
E86
E87
E88
E89
E90
E91
E92
E93
E94
E95
E96
L1
L2
L3
L4
L5
L6
L7
L8
L9
L10
L11
L12
L13
L14
L15
L16
L17
L18
L19
L20
L21
L22
L23
L24
L25
L26
L27
L28
L29
L30
L31
L32
L33
L34
L35
L36
L37
L38
L39
L40
L41
L42
L43
L44
L45
L46
L47
L48
L49
L50
L51
L52
L53
L54
L55
L56
L57
L58
L59
L60
L61
L62
L63
L64
L65
L66
L67
L68
L69
L70
L71
L72
L73
L74
L75
L76
L77
L78
L79
L80
L81
L82
L83
L84
L85
L86
L87
L88
L89
L90
L91
L92
L93
L94
L95
L96
RPKM
0
25
50
B
XIST
***
log2 RPKM
5
0
-5
-10
Early naïve
Late naïve

**Figure S1, related to figure 1:** Transition of early naïve to late naïve state is associated with increased *XIST* expression. (A) Heatmap representing the *XIST* expression (RPKM) in early naïve (n=96 cells) and late naïve cells (n=96 cells). (B) Comparison of *XIST* expression between early naïve (n=96 cells) and late naïve cells (n=96 cells). $p<0.00001$ (Mann-Whitney U-test).

Fig. S2

**A**

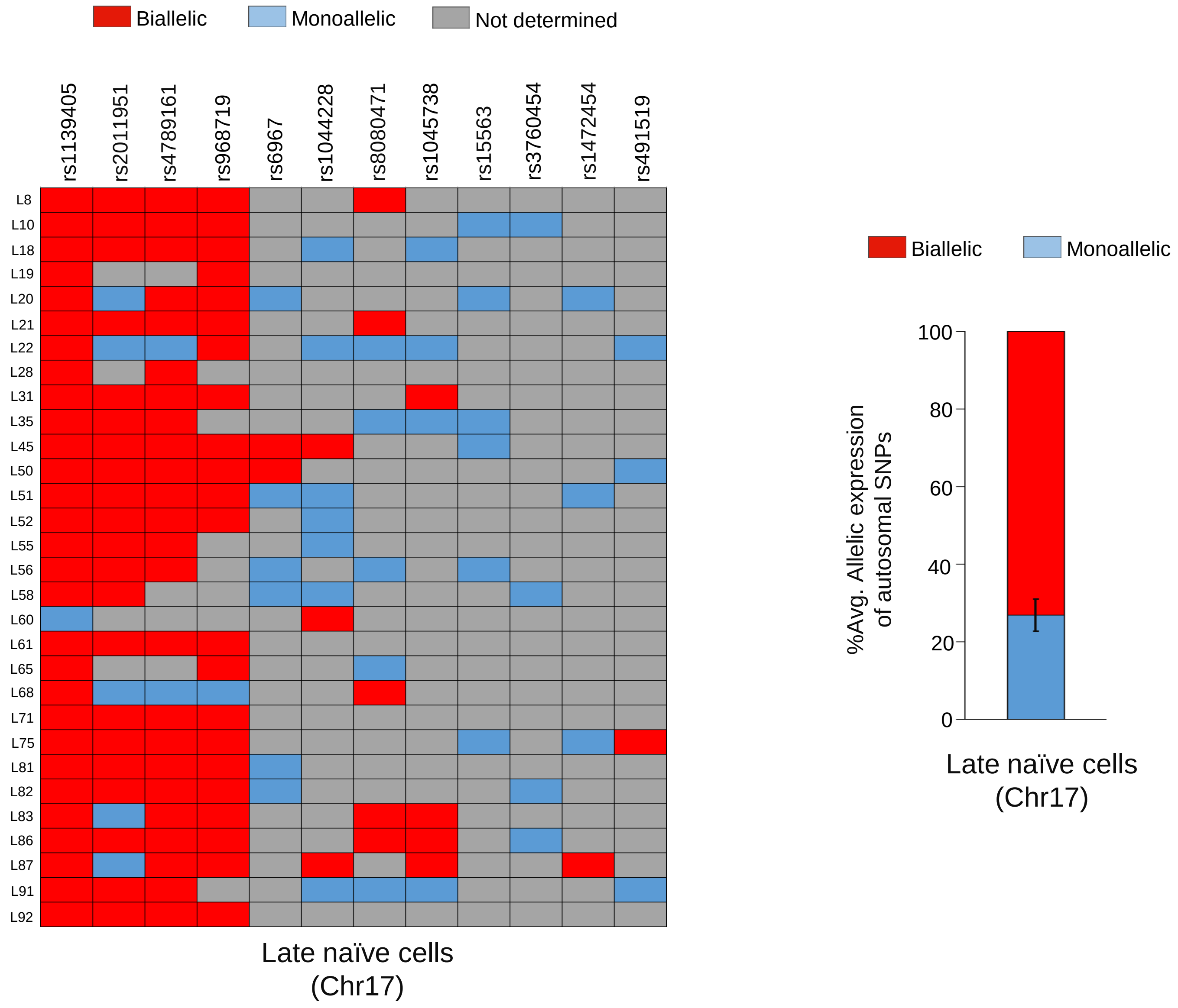
Biallelic
Monoallelic
Not determined
rs1139405
rs2011951
rs4789161
rs968719
rs6967
rs1044228
rs8080471
rs1045738
rs15563
rs3760454
rs1472454
rs491519
L8
L10
L18
L19
L20
L21
L22
L28
L31
L35
L45
L50
L51
L52
L55
L56
L58
L60
L61
L65
L68
L71
L75
L81
L82
L83
L86
L87
L91
L92
Late naïve cells (Chr17)
Biallelic
Monoallelic
%Avg. Allelic expression of autosomal SNPs
0
20
40
60
80
100
Late naïve cells (Chr17)

**B**

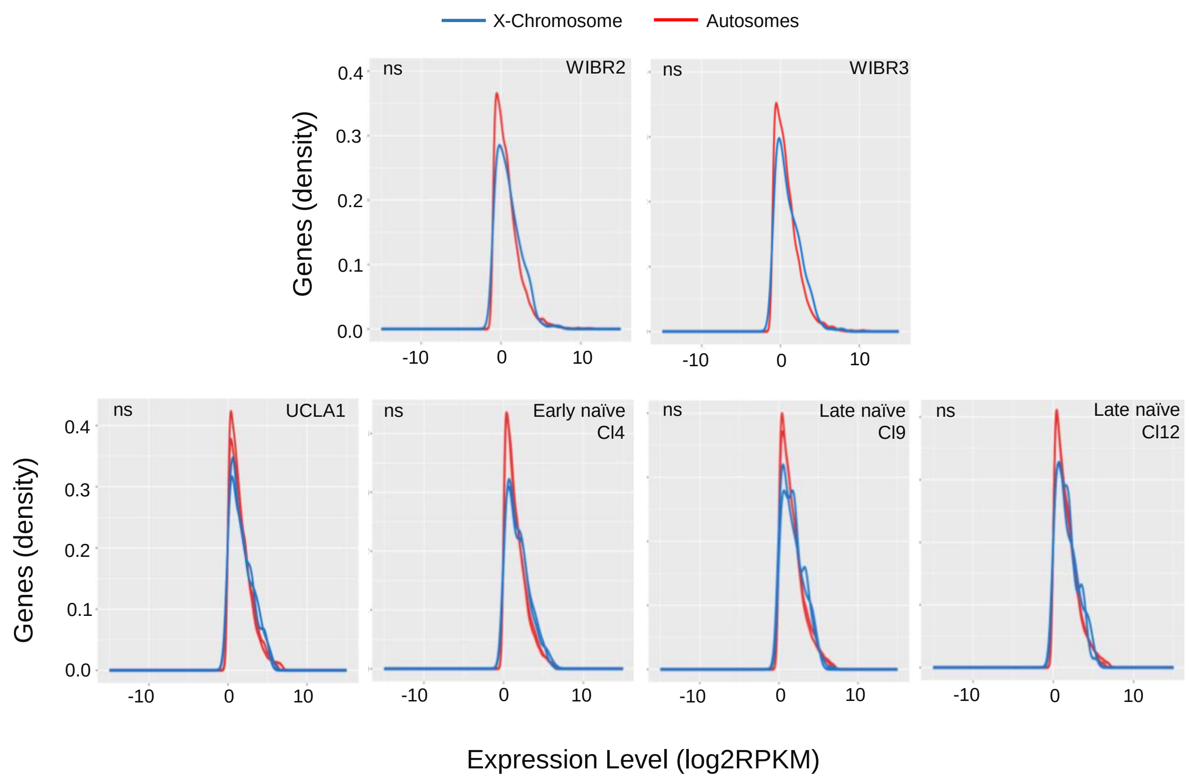
X-Chromosome
Autosomes
ns
WIBR2
ns
WIBR3
Genes (density)
0.0
0.1
0.2
0.3
0.4
-10
0
10
ns
UCLA1
ns
Early naïve Cl4
ns
Late naïve Cl9
ns
Late naïve Cl12
Genes (density)
Expression Level (log2RPKM)

**Figure S2, related to figure 3 and figure 4:** (A) Allelic expression analysis of autosomal genes (Chr17) in late naïve cells. Right, histogram showing the quantification of average percent of SNPs showing monoallelic and biallelic in late naïve cells. (B) Histograms showing that there were no significant difference between X-linked and autosomal gene expression distribution for WIBR2, WIBR3, UCLA1 primed, UCLA1 early naïve and UCLA1late naïve cells. ($p > 0.05$, by Kolmogorov-Smirnov test).

**Supplementary file legends**

**Supplementary file 1, related to Figure 1:** Analysis of X-linked gene expression in early and late naïve cells.

**Supplementary file 2, related to Figure 2:** Allelic analysis of *XIST* expression and comparison of X-linked gene expression in *XIST*-monoallelic, -biallelic and -negative cells of late naïve state.

**Supplementary file 3, related to Figure 3:** Allelic analysis of X-linked genes and autosomal genes (Chr17) in early and late naïve cells.

**Supplementary file 4, related to Figure 3:** Analysis of X-chromosomal ploidy in early and late naïve cells.

**Supplementary file 5, related to Figure 4:** RPKM of X-linked and autosomal genes used for X:A ratio analysis.

**Supplemental experimental procedures**

**Data acquisition:** For active X-chromosome upregulation analysis (related to Fig. 4), we used following datasets from bulk-population RNA-Seq: hESCs (H1) / 3iL hESCs (H1) -E-MTAB-2031(Chan et al., 2013), WIBR1, WIN1, WIBR2, WIBR3- GSE75868 (Theunissen et al., 2016), WIS2-GSE60138 (Irie et al., 2015), UCLA2- GSE88933 (Patel et al., 2017), H1_A-GSE85331 (Liu et al., 2017), HUES8 - GSE102311(Sun et al., 2018), HNES1_A, HNES1_B - E-MTAB-5674 (Guo et al., 2017), HNES1_C-E-MTAB-4461(Guo et al., 2016), 4iLIF H1, Lis1-GSE60955 (Sperber et al., 2015), H1_B-GSE75748 (Chu et al., 2016). For hESCs (Sc) - GSE36552 (Yan et al., 2013), we used single cell RNA-Seq dataset of 8 single cells.

**Origin of naïve hPSCs:** WIN1: Embryo derived, cultured in either 5i/L/A or 4i/L/A medium (Theunissen et al., 2016); 3iL hESCs: converted from hESCs (H1) using 3iL media (Chan et al., 2013); HNES1_A / B / C: Embryo derived, cultured in t2iLGO in different feeder conditions (Guo et al., 2016, 2017); 4iLIF H1: converted from H1 using 4iLIF media (Sperber et al., 2015); NHSM Lis1: Embryo derived, cultured in NHSM media (Sperber et al., 2015).

**Reads mapping and counting:** Reads were mapped to the human genome (hg38) using STAR (Dobin et al., 2013). Mapped reads were then processed using SAM tools (Li et al., 2009) and the number of reads mapping to each gene was counted using HTSeq-count (Anders et al., 2015).

**Expression analysis of *XIST* and X-linked genes in early vs late naïve cells:** We calculated RPKM (Reads Per Kilobase per Million) using rpkmforgenes (Ramsköld et al., 2009). First, we checked *XIST* expression in 192 single cells consisting of 96 early naïve and 96 late naive cells. Then, for early naive cells we selected 60 cells out of 96 showing very low *XIST* expression (0 to 0.5 RPKM) and then from these 60 cells we selected top 28 early cells having RPKM sum (2634.96 to 1267.39) and mean (> 0.9) for further analysis. For late naive cells, we selected 59 out of 96 showing very high *XIST* expression (3 to 39 RPKM) and then from these 59 cells we selected top 30 cells having RPKM sum (1949.62 to 1195.90) and mean (> 0.9) for further analysis. To check the expression of X-linked genes in (28 early naive + 30 late naive cells), we considered only those genes which were having $5 < \mu_{RPKM} < 200$.

**Expression analysis of X-linked genes in *XIST*-biallelic, -monoallelic and -negative late naïve cells:** For this analysis, 59 cells were selected out of 96 late naive cells based on the higher *XIST* expression (3 - 39 RPKM). Out of these 59 cells, 35 cells (RPKM sum >500; RPKM mean > 0.4) had allelic data for *XIST*. Allelic expression analysis for *XIST* was performed as described in allelic expression analysis method section. Out of these 35 cells, 26 cells having monoallelic expression and 9 cells having biallelic expression for *XIST* were chosen for X-linked gene expression analysis. 17 *XIST* negative cells were selected based on *XIST* RPKM<1 and having RPKM sum >500; RPKM mean > 0.4. X-linked genes for expression analysis were chosen based on the $5 < \mu_{RPKM} < 200$.

**Ploidy:** To analyze the X-chromosomal ploidy, we considered 70 genes having RPKM mean ≥ 3 distributed across the X-chromosome for 58 cells (28 early naive cells + 30 late naive cells). We calculated gene expression ratio through dividing the individual gene's RPKM by its median value across all the 58 cells. To visualize the profile of X-chromosome in all these cells, the normalised gene expression values (gene expression ratio) were used to build a moving average plot (neighbourhood size, k=9) using methods described by Mayshar *et al.* (Mayshar et al., 2010).

**X-chromosome to autosomes expression ratio:** We calculated X:A ratio by dividing the median expression (RPKM) of the X-linked genes by the median expression (RPKM) of autosomal genes. For this analysis we removed the no/low expressed genes and considered the genes having ≥ 0.5 RPKM for both X and autosomal genes. We decided this threshold, based on the previous reports showing that exclusion of low expressed genes below 0.5 FPKM are more appropriate for assaying the active-X chromosome upregulation (Deng et al., 2011; Li et al., 2017; Sangrithi et al., 2017; Yildirim et al., 2012). We also profiled the expression level distribution of X-linked and autosomal genes as density plots created in R using the ggplot2 package. However, for analysis of X:A ratio of UCLA1 primed, early naïve and late naïve cells, we considered the expressed genes with RPKM ≥1 and upper RPKM threshold that corresponded to the lowest 99th centile RPKM value of expression to avoid the difference between X-linked and autosomal gene expression distribution. We excluded escapees and genes in the pseudo autosomal regions of X-chromosome for our analysis.

**Statistical tests and Box-plots:** All statistical tests and boxplots were performed using R version 3.5.1(R Development Core Team, 2016).
